# Association between Japanese community health workers’ willingness to continue service and two categories of motives: altruistic and self-oriented

**DOI:** 10.1101/703322

**Authors:** Atsuko Taguchi, Hiroshi Murayama, Keiko Ono

## Abstract

**Background:** As population aging progresses, volunteers in health field are expected to play a key role in health promotion and disease prevention, which may improve community residents’ health and well-being and at the same time help curve healthcare cost. The objective of this study is to examine the effects of self-oriented motives and altruistic motives as explanatory factors for Japanese Community Health Workers (CHWs)’ desire to continue their service. Unraveling the relative effects of these two types of motivation on CHW retention may lead to policy and practical implications for recruiting, training, and supporting CHWs in Japan. Haddad (2007) observed that citizens in Japan generally have a sense of governmental and individual responsibility for dealing with social problems. Applying these insights to CHWs, we hypothesize that altruistic motives have more potent influence on volunteers’ willingness to continue to serve than self-oriented motives.

**Methods:** Three cities in Shiga prefecture, Japan agreed to participate in the study. Anonymous, self-administered questionnaire was mailed to all CHWs who work in the three communities. The survey data were collected in March and April, 2013. A total of 417 questionnaires were mailed to CHWs, of which 346 were completed and returned (response rate 83.0%). Nine questionnaires missing response to the question concerning willingness to continue serving were removed from the analysis. The final analysis used 337 questionnaires (effective response rate 80.8%).

**Results:** One hundred ninety-nine (59.1%) of the respondents answered the question about willingness to continue CHW affirmatively, and 138 (40.9%) negatively. Controlling for other relevant factors, those with self-oriented motives in serving as CHWs were more likely to state they are willing to continue to serve (OR:1.54, confidence interval 1.00-2.37) than those without such motives. Contrary to our hypothesis, the effect of altruistic motives on respondents’ willingness to continue serving as CHWs was slightly larger than that of self-oriented motives; the odds ratio was 1.56(confidence interval 1.08-2.27).

**Conclusion:** One practical implication of the research is that learning more about the twin motives, self-oriented and altruistic, of volunteers and tailoring the content of CHW training to address those motives may be beneficial.

## Introduction

Though population aging can be observed in virtually all regions of the world today [1], its advancement is most pronounced in the affluent nations. The rapidly aging population in such countries has led to a number of policy challenges, one of the most important of which is an increase in healthcare costs [2]. To help contain such cost increase, there has been a growing emphasis on health promotion and disease prevention to fight chronic, lifestyle related diseases [3–4] and to help older adults maintain their independence [5]. Health promotion and disease prevention measures targeting the elderly have been shown to not only mitigate the risk of common health issues such as falls but also to improve socio-psychological well-being of the recipients [6].

It is increasingly difficult, however, for the government in industrialized nations to meet the needs of all of its citizenry given the limited resources [7], and health promotion and prevention is no exception. In response to such challenges, there is a growing interest in utilizing *volunteers* to provide health promotion and education services as a way to help fill the gap between the population’s needs and the resources available [7–8]. There is already an established tradition of community health workers (CHWs), most of whom are uncompensated volunteers, in many nations [9]. CHWs engage in health promotion and disease prevention by connecting residents to appropriate and available medical care in the community, encouraging residents to use services, and facilitating access [9]. Community health workers delivering health information and advice to their neighbors could prove to be one of the keys to maintaining the physical as well as financial health of the nation [9].

Japan is among the most advanced aged (and aging) society in the world; its proportion of older adults (age 65 and older) was the highest in the world (27.1%; Li et al, 2019) [1] in 2017 and is expected to reach 40.0% in 2060 [10]. Consequently, health promotion volunteers in Japan, one variant of CHWs, increasingly focus on both supporting the health needs of the elderly and educating all generations about prevention of disease and disability that would manifest later in one’s life [11]. Though the tradition of CHWs in Japan dates back to around 1930, the CHW priorities have changed drastically since then as society’s health needs have changed; the initial emphasis on prevention of infectious disease and improving nutrition was gradually replaced by efforts to prevent chronic diseases associated with aging and to stave off frailty among the elderly. Roughly 80 percent of the Japanese municipalities today recruit, train, and commission such CHWs [12] who provide counseling, education about various health concerns, and mobilization of community members and resources.

One major challenge in utilizing volunteers including CHWs in the social service and health sector is recruitment and retention [13–14]. The total number of volunteers serving in hospitals, for instance, is increasing, but at the same time “premature dropping out” is also on the rise [13]; the high turnover means a higher recruitment and training cost and possible service disruption for the organizations [13,15] as well as missed opportunity to reap full physical and other benefits from participation for the volunteers [16]. In the Japanese context, the overall decline in volunteering as CHWs has been attributed to the rising level of female labor force participation as CHWs are predominantly home makers [17]. In addition, as the household size decreases, family members are responsible for a greater share of the child care and elderly care duties which leaves less time for volunteering as a CHW [18]. Reflecting such changes in society, two additional health related volunteer institutions in Japan have seen large decline in their ranks; the number of healthy eating promotion volunteers has been decreasing steadily since its peak in 1998 (220,000) [19], and maternal health support volunteers went from 70,000 (1993) to 42,000 in 2013 [20].

Davis and associates present an elaborate model of how certain antecedents including altruistic and self-oriented motivations contribute to volunteers’ decision to participate in the first place, and subsequently how volunteers’ experiences mediate further involvement and continuation of services [21]. The model presented suggests that fulfilling volunteers’ motivations increases their satisfaction level and thereby encourages their continued participation. Applying this insight to CHWs, understanding how different motivations influence health volunteers to persist in their activities will help promote volunteers’ active involvement and retention. Studies on volunteer retention have found ambiguous effects of motivation on volunteers’ desire to continue serving [14], and moreover, there is dearth of studies that examine CHWs’ motivation to continue their service beyond the initial period. Specifically in Japan, we are not aware of any empirical study explaining CHW retention that focuses on volunteers’ motivation. As a survey of municipalities in Japan has found, many communities report that short tenure of volunteers is among the challenges [12]. Identifying motivational and other factors that contribute to CHWs’ willingness to continue to serve will help inform measures that can be taken to maintain volunteer morale and thus promoting retention and additional recruitment [22].

Psychological studies on volunteer motivation have identified two broad categories of motivations for volunteer participation: self-oriented motives and altruistic motives [21–23]. Self-oriented motives for volunteering include desire for personal development and learning, and altruistic motives include desire to help others and to fulfill social responsibility [22]. Initially, research focused on the altruistic motives as the key ingredient for volunteer participation [22, 24]; more recently, the dominant approach has been one that assumes that motives for volunteer participation are multi-faceted, contextual, and complex [21, 23–25]. In the Japanese context, Sakurai [26] investigated the association between volunteer motives for participation and the characteristics of both the target activities and the volunteers such as age and gender; the findings show that altruistic motives had greater influence than self-oriented motives on older volunteers’ willingness to continue service.

The objective of this study therefore is to examine the effect of self-oriented motives and altruistic motives [22] as explanatory factors for Japanese CHWs’ desire to continue their service. Unraveling the relative effects of these two types of motivation on CHW retention may lead to policy and practical implications for recruiting, training, and supporting CHWs in Japan. Haddad [27] observed that citizens in Japan generally have a sense of governmental and individual responsibility for dealing with social problems. Applying these insights to CHWs, we hypothesize that altruistic motives have more potent influence on volunteers’ willingness to continue to serve than self-oriented motives.

## Data and Methods

### Research areas and participants

The research was conducted in three communities (*Kusatsu* city, *Ritto* city, *Yasu* city) in southern part of *Shiga* prefecture, Japan. Southern *Shiga* is a bedroom community which is half an hour train ride away from the city of Kyoto. The total population in each city was 137,000 (*Kusatsu* city), 67,000 (*Ritto* city), and 50,000 (*Yasu* city) as of 2016. The proportion of elderly (age 65 and older) was lower than the national average: 19.6% in *Kusatsu* city, 17.6% in *Ritto* city, and 24.0% in *Yasu* city.

To be commissioned as a CHW, one must complete at least 20 hours of training. The public health center located in each city organizes and offers such training, and these centers also offer additional training and support for the active CHWs. Each city runs its own CHW program, yet, the programs are quite similar across communities. The total number of CHWs at the time of the research was as follows: 180 (*Kusatsu* city), 157 (*Ritto* city), 80 (*Yasu* city).

### Methods

Anonymous, self-administered questionnaire was mailed to the CHWs. The survey data were collected in March and April, 2013. Initially, the research team met with public health center officials in the three cities to explain the objective of the study and to request their assistance in the distribution of the survey; subsequently, the questionnaire was sent to all 417 CHWs serving in the three cities which agreed to participate in the study.

This study was approved by the ethics review board of the Tokyo Metropolitan Institute of Gerontology. The research team explained the objective of the research to the CHWs, city and public health officials in the three cities which then gave their consent to take part in the study. A letter was attached to each mailed questionnaire; the letter explained the purpose of the survey and clearly stated that participation was voluntary, and data confidentiality will be protected. Filling out and sending back the survey constituted consent.

### Measures

#### Willingness to continue to serve as CHW

CHWs’ desire to continue serving was measured with a single Likert scale item; the participants were asked whether they agree or not with the statement “I would like to continue to serve as a CHW” (1: agree, 2: somewhat agree, 3: somewhat disagree, 4: disagree). For ease of interpretation, the scale variable was converted into a dummy variable (1 if agree or somewhat agree, 0 if disagree or somewhat disagree),

#### Altruistic and self-oriented motives for serving as CHWs

The motivational factors related to who gets and stays involved that researchers have focused on are fulfillment of altruistic and self-oriented motives [7, 21–24, 28]. Self-oriented motives include acquisition of new knowledge and skills while altruistic motives include desire to help others and to contribute to society [7]. The existing studies have found some support for both self-oriented and altruistic motives as the drivers for volunteer retention, and several studies in particular report findings supporting the role of self-oriented motives [7, 29–30]. One study conducted in Australia found that other-oriented motives were positively correlated while self-oriented motives were *negatively* correlated with intentions to continue volunteering [28].

To measure the respondents’ self-oriented and altruistic motives in serving as CHWs, the survey also asked if they agreed with the following statements: “I do understand and appreciate the benefits of serving as an CHW for myself (such as gaining knowledge about health)” (self-oriented), “I do understand and appreciate the benefits of CHW activities for community residents (such as CHW being able to offer assistance in informal setting)” (altruistic). The questions were constructed as six-point scales with responses ranging from 1: strongly disagree to 6: strongly agree.

#### Individual characteristics

Existing scholarly research has examined who initiates and then persists in volunteer activities [14]. The demographic characteristics which have been shown to have an effect on volunteer participation include age, sex, occupation, and race [7, 30]. The questionnaire also asked respondent’s age, sex, educational attainment, work status, household economic situation, self-rated health, and household composition. The educational attainment had three response categories: high school graduates (or less), junior college and other two-year tertiary educational institution graduates, four-year college graduates.

Work status question had several response categories which include full time and part time employment. The work status variable was converted into a dummy variable (1 if working full or part time or self-employed, 0 if not working. Full time homemaker was coded 0). As for household economic situation, respondents were asked to choose from 1: barely paying bills to 5: comfortable. Self-rated health is a four-point scale (1: not healthy, 4: healthy).

#### Factors that may explain CHW motivation

The respondents’ years of service as CHW, their level of motivation at the completion of initial training, and whether they have served in a volunteer leadership role were also assessed. The level of motivation was asked by a six-point scale question (1: not at all motivated, 6: highly motivated).

### Analytical Methods

First, bivariate analysis was conducted to examine the relationships between the CHWs’ willingness to continue to serve and each of the factors that may explain such willingness identified in the literature. T test was applied to numeric variables, and Chi square test was conducted on categorical variables. Next, multivariate logistic regression analysis was conducted with the volunteers’ desire to continue serving as the dependent variable (1 if willing to continue to serve, 0 if not). The main independent variables of interest are the self-oriented motive (understand and appreciate the benefits of serving to oneself) and the altruistic motive (understand and appreciate the benefits to local residents). Individual characteristics and a few additional variables found to be relevant in the existing literature are added as control variables.

## Results

### Response rate and descriptive characteristics of the research participants

A total of 417 questionnaires were mailed to CHWs, of which 346 were completed and returned (response rate 83.0%). Nine questionnaires missing response to the question concerning willingness to continue serving were removed from the analysis. The final analysis used 337 questionnaires (effective response rate 80.8%). One hundred ninety-nine (59.1%) of the respondents answered the question about willingness to continue CHW affirmatively, and 138 (40.9%) negatively. The descriptive characteristics of the respondents are shown in Table 1. The average age of the volunteers was 62.2 years (standard deviation 7.2), and an overwhelming majority were women (330 or 97.9%). The most common educational attainment was completion of middle school or high school (n=216, 64.1%), followed by completion of two-year tertiary education institution (n=97, 28.8%), then completion of four-year college (n=20, 5.9%). A majority (n=200, 59.3%) were not employed. The mean self-rated health score was 3.2 (1: not healthy, 4: healthy) with standard deviation 0.7. Only a very small fraction of the volunteers (n=12, 3.6%) lived by themselves; the remaining 320 (95%) lived with family member(s). The mean number of years of service as CHW was 8.5 years (standard deviation 6.0).

**Table 1.** Characteristics of the research participants

|  | Total<br>n=337<br>n (%) | Willingness to continue<br>to serve as CHW |  | p-value |
| --- | --- | --- | --- | --- |
|  |  | No<br>n=138<br>n (%) | Yes<br>n=199<br>n (%) |  |
| **Age**<sup>1)</sup> | 62.2±7.2 | 61.3±7.4 | 62.4±7.0 | 0.168 |
| **Sex** |  |  |  |  |
| Men | 7 (2.1) | 2 (1.4) | 5 (2.5) | 0.705a |
| Women | 330 (97.9) | 136 (98.6) | 194 (97.5) |  |
| **Educational attainment** |  |  |  |  |
| High school graduates or less | 216 (64.1) | 89 (64.5) | 127 (63.8) | 0.861 |
| Junior college or other two-year tertiary educational institution graduate | 97 (28.8) | 39 (28.3) | 58 (29.1) |  |
| Four-year college graduates | 20 (5.9) | 7 (5.1) | 13 (6.5) |  |
| **Work status** |  |  |  |  |
| Working | 200 (59.3) | 76 (55.1) | 124 (62.3) | 0.238 |
| Not working | 133 (39.5) | 76 (55.1) | 73 (36.7) |  |
| **Household economic situation** [range: 1-5] | 3.5±0.8 | 2.9±0.8 | 2.7±0.8 | 0.012 |
| **Self-rated health**<sup>1)</sup> [range: 1-4] | 3.2±0.7 | 3.0±0.7 | 3.3±0.6 | <0.001 |
| **Household composition** |  |  |  |  |
| Living alone | 12 (3.6) | 3 (2.2) | 9 (4.5) | 0.373a |
| Living together with family member(s) | 320 (95.0) | 131 (94.9) | 189 (95.0) |  |
| **Years of service as CHW**<sup>1)</sup> | 8.5±6.0 | 8.0±5.7 | 8.9±6.2 | 0.199 |
| **Level of motivation at the completion of initial training**<sup>1)</sup> [range: 1-6] | 4.2±0.9 | 3.8±1.0 | 4.5±0.8 | <0.001 |
| **Experiences of CHW leadership role** |  |  |  |  |
| No experiences | 169 (50.1) | 80 (58.0) | 89 (64.5) | 0.018 |
| Experiences | 162 (48.1) | 56 (40.6) | 106 (76.8) |  |
| **Self-oriented motives in serving as CHW**<sup>1)</sup> [range: 1-6] | 4.2±0.9 | 3.9±0.9 | 4.4±0.7 | <0.001 |
| **Altruistic motives in serving as CHW**<sup>1)</sup> [range: 1-6] | 3.5±1.0 | 3.2±1.0 | 3.7±0.9 | <0.001 |

<sup>1)</sup> Mean±Standard deviation
a: Fisher's exact test
no mark: Chi-square test

### Altruistic and self-oriented motives analysis

Controlling for other relevant factors, those with self-oriented motives in serving as CHWs were more likely to state they are willing to continue to serve (OR: 1.54, confidence interval 1.00-2.37) than those without such motives. Contrary to our hypothesis, the effect of altruistic motives on respondents’ willingness to continue serving as CHWs was slightly larger than that of self-oriented motives; the odds ratio was 1.56(confidence interval 1.08-2.27) (Table 2).

**Table 2.**
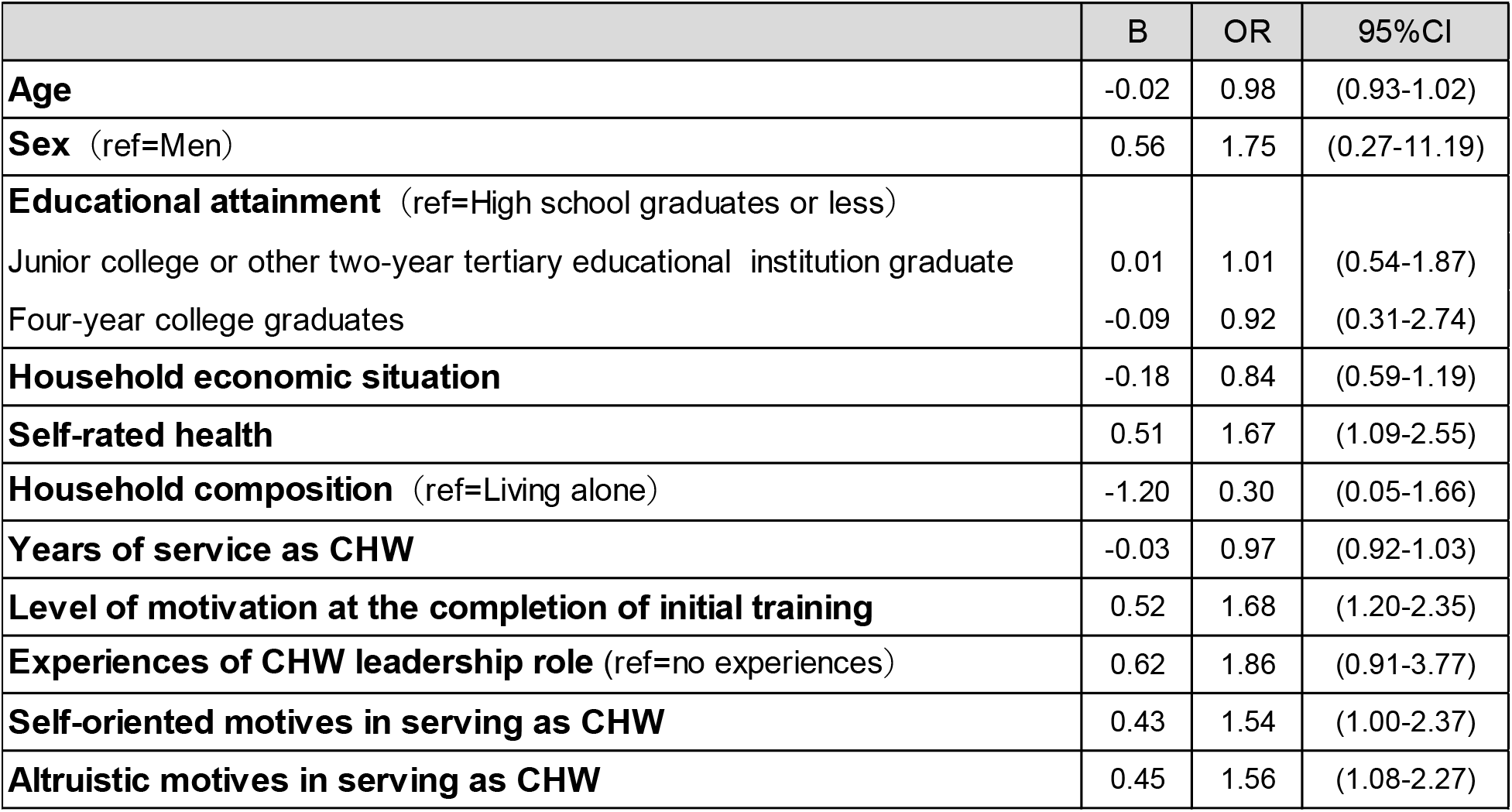
Multivariate analysis for willingness to continue to serve as CHW

Both self-rated health (OR: 1.67, confidence interval 1.09 – 2.55) and the volunteer’s level of motivation at the completion of initial training (OR: 1.68, confidence interval 1.20 – 2.35) had positive and statistically significant effects on the willingness to continue to serve as hypothesized. There was no statistically significant association between desire to continue volunteering as CHW and age, sex, educational attainment, household economic situation, year of service as CHW, and experiences of CHW leadership role.

## Discussion

### The characteristics of the sample

Approximately 60% of the respondents expressed willingness to continue serving as CHWs. The sample was virtually all female (97.9 %) and the average age was 62.2 years old which is significantly higher than the median age in these communities. The majority of the sample (59.3%) were not working at the time of the survey (“not working” includes full time homemakers). Such characteristics of the sample indicates that the sample represents CHWs in Japan well; a national survey conducted earlier in Japan [12] has found that CHWs tend to be majority female and older. There is evidence that suggests women tend to be more interested and engaged in health issues than men, and age has a positive relationship with interest in health [31]. Previous studies have also found that women greatly outnumber men among health volunteers, and the volunteers tend to be older [7, 13, 22].

### Willingness to continue serving and self-oriented and altruistic motives

In this study, controlling for other relevant factors, those who understand and appreciate the benefits of serving as CHW to oneself were more likely to state they are willing to continue to serve (OR:1.54, confidence interval 1.00-2.37). Our findings confirm what several earlier studies [24, 30, 32] have found; both self-oriented motives and altruistic motives were related to the volunteers’ willingness to continue in their capacity. Haddad [27], makes the case that many citizens in Japan volunteer in activities that have “close, embedded relationship with the local government.” [27]; in her findings, those who believe the government, rather than individuals, must deal with social issues and those who consider volunteering to be civic duty are more likely to serve. Furthermore, Sakurai [26] found that homemakers and older adults who volunteer in Japan are motivated by altruism. In the present research, we hypothesized that altruistic motives have a more robust impact on the CHWs’ retention than self-oriented motives in Japanese CHW organizations characterized by deep local ties; the empirical analysis, however, showed that self-oriented motives work just as strongly as altruistic motives. This might be explained by the transformation of the Japanese society in the last few decades; information technology has been deeply integrated to all corners of Japan which provides higher quality of life on one hand, but on the other hand has resulted in local community organizations becoming less relevant [18]. Haddad [27] observed that citizens in Japan have a sense of governmental and individual responsibility for dealing with social problems. Our findings may indicate that Japanese citizens’ sense of community responsibility may in fact have been waning. Our confirmation that self-oriented motives also affect CHWs motivation to continue serving in a local community organization is a significant contribution to the literature on CHW retention.

### Practical Implications

The findings of this research can be applied to inform CHW training programs. In the countries where CHW recruitment and training has been studied extensively, researchers have concluded that CHW training programs generally lack consistency in both length and curriculum content across administrative units [9]. In this context, the content of the CHW training program has rarely been evaluated from the perspective of self-oriented and altruistic motives.

In Japan, the most important CHW activities include encouraging local residents to get health check-up and conducting health education classes. In CHW training, volunteers acquire health information and knowledge, but the emphasis has primarily been on the information needs of local residents, not the volunteers. This research indicates that, in addition to altruistic motives, self-oriented motives play a significant role in volunteer retention. One practical implication of the research is that learning more about the twin motives, self-oriented and altruistic, of volunteers and tailoring the content of CHW training to address those motives may be beneficial. For instance, many CHWs in Japan are older adults, which means that they often have health issues and concern of their own; providing volunteers with training that addresses volunteers’ health needs and opportunities to attend health education classes may be successful in ensuring continued service.

### Limitations

Our research design is cross-sectional. To observe whether the volunteers actually continue their health volunteer activities or not, one needs to employ a longitudinal research design. In the survey, because of space limitation, a single question was used to assess the volunteers’ self-oriented and altruistic motives. Further research is needed to evaluate and improve the reliability and validity of the question(s) and the scale. One additional limitation of the present research is it involved a small number (n=3) of health volunteer organizations located in the same region of a single prefecture in Japan.

## Conclusions

This study explored the effects of self-oriented and altruistic motives on community health volunteers’ willingness to continue serving in Japan. Controlling for other relevant factors, those with self-oriented motives in serving as CHWs were more likely to state they are willing to continue to serve (OR:1.54, confidence interval 1.00-2.37) than those without such motives. Contrary to our hypothesis, the effect of altruistic motives on respondents’ willingness to continue serving as CHWs was slightly larger than that of self-oriented motives; the odds ratio was 1.56(confidence interval 1.08-2.27).

One practical implication of the research is that learning more about the twin motives, self-oriented and altruistic, of volunteers and tailoring the content of CHW training to address those motives may be beneficial.

## Acknowledgments

This research (“Verification of the effects of resident organization led outreach model to prevent social isolation”) was conducted with grants from the Pfizer Health Research Foundation. The principal investigator was Atsuko Taguchi.

## Supporting information

Data set

